# Using Bayesian Inference to Estimate Plausible Muscle Forces in Musculoskeletal Models

**DOI:** 10.1101/2021.07.28.454251

**Authors:** Russell T. Johnson, Daniel Lakeland, James M. Finley

## Abstract

**Background:** Musculoskeletal modeling is currently a preferred method for estimating the muscle forces that underlie observed movements. However, these estimates are sensitive to a variety of assumptions and uncertainties, which creates difficulty when trying to interpret the muscle forces from musculoskeletal simulations. Here, we describe an approach that uses Bayesian inference to identify plausible ranges of muscle forces for a simple motion while representing uncertainty in the measurement of the motion and the objective function used to solve the muscle redundancy problem.

**Methods:** We generated a reference elbow flexion-extension motion by simulating a set of muscle excitation signals derived from the computed muscle control tool built into OpenSim. We then used a Markov Chain Monte Carlo (MCMC) algorithm to sample from a posterior probability distribution of muscle excitations that would result in the reference elbow motion trajectory. We constructed a prior over the excitation parameters which down-weighted regions of the parameter space with greater muscle excitations. We used muscle excitations to find the corresponding kinematics using OpenSim, where the error in position and velocity trajectories (likelihood function) was combined with the sum of the cubed muscle excitations integrated over time (prior function) to compute the posterior probability density.

**Results:** We evaluated the muscle forces that resulted from the set of excitations that were visited in the MCMC chain (five parallel chains, 450,000 iterations per chain, runtime = 71 hours). The estimated muscle forces compared favorably with the reference motion from computed muscle control, while the elbow angle and velocity from MCMC matched closely with the reference with an average RMSE for angle and velocity equal to 0.008° and 0.18°/s, respectively. However, our rank plot analysis and potential scale reduction statistics, which we used to evaluate convergence of the algorithm, indicated that the parallel chains did not fully mix.

**Conclusions:** While the results from this process are a promising step towards characterizing uncertainty in muscle force estimation, the computational time required to search the solution space with, and the lack of MCMC convergence indicates that further developments in MCMC algorithms are necessary for this process to become feasible for larger-scale models.

## Background

Movement scientists are often interested in quantifying the timing and magnitude of muscle forces during motions like walking or reaching to understand causal links between muscle mechanics and movement. Accurately and reliably estimating individual muscle forces has implications for how well researchers can evaluate muscle function to help guide surgical interventions, inform the design of prosthetics and orthotics, and estimate other clinically relevant outputs (e.g., joint contact forces) (1–6). Measuring muscle forces *in vivo* is difficult to do, except on a limited scale (e.g., triceps surae forces (7)), but most often the methodology is far too invasive to use with human participants. Instead, researchers often use experimental data combined with musculoskeletal modeling to estimate muscle forces during a movement (8–11). Several methods have been developed to estimate individual muscle forces during a motion, including static optimization, computed muscle control, and electromyography (EMG) optimization (12–14). Typically, these methods result in a single force trajectory for each muscle that optimizes a chosen objective function for a given musculoskeletal model and experimental motion. However, accurately estimating muscle forces remains difficult because the musculoskeletal system is redundant (an infinite combination of muscle forces can often give rise to the same joint moment) (15), and simulations of movement depend on experimental data and a variety of parameters that are prone to uncertainties (16,17).

The uncertainty associated with muscle force estimation can arise from the uncertainty with which we mathematically represent how the central nervous system distributes muscle forces amongst agonist muscles (8,12), errors in marker placement and skin movement relative to anatomical landmarks (18,19), and modeling assumptions related to muscle parameters (20,21). Typical representations of motor control assume that the central nervous system attempts to minimize some objective function (e.g., minimize muscle fatigue or metabolic energy cost (8,16,22)). Choosing an appropriate objective function for a particular motion is difficult because it is not known exactly how the nervous system distributes forces across muscles. In reality, the nervous system is unlikely to generate an “optimal” motion under any hypothesis represented by a simple objective function. The unknowns associated with choosing an appropriate objective function for musculoskeletal simulations are problematic because muscle force estimates are sensitive to the objective function chosen (16,22–25). Other aspects of musculoskeletal modeling that can lead to uncertainty in the accuracy of estimated muscle forces such as variability in marker placement, movement artifact, and unknown model parameters can also play a role in impacting the computed muscle forces (17–20,26–28). With uncertainties in the motor control model, the measured data, and the musculoskeletal model, it is clear that the single solutions typically obtained from standard static or dynamic optimizations conceal the inherent uncertainty we have about the predicted muscle forces.

One approach to quantifying uncertainty in musculoskeletal modeling is to treat the objective function as unknown while keeping the other model parameters fixed. There have been a few different methods developed to try to capture some of the uncertainty associated with choosing an objective function and how it would affect the estimated muscle forces for a given model or motion (29–31). These approaches have used mathematical mappings between joint torques and muscle activations to compute upper and lower bounds on the muscle forces for each muscle over time. Instead of choosing an explicit objective function to solve the muscle redundancy problem, these methods instead solve for the upper and lower bounds then assume the muscle forces lie somewhere in between. However, these ranges include solutions that would only be possible with extreme co-activation of agonist and antagonist muscles. Extreme co-activation is unlikely to occur in most healthy human sub-maximal movements, especially if muscle forces are distributed in a way that is sensitive to the physiological load (or effort) across individual muscles (22) or reduces metabolic cost. One previous study used EMG data as a way to provide some bounds on the range of possible muscle forces (29), however the remaining muscles without EMG data were left unbounded and therefore still had vast ranges of possible muscle forces. Additionally, there are other limitations to directly using EMG data for this approach, such as uncertainty about how to normalize EMG, resolving forces from EMG, and collecting EMG from deep muscles (32–35). Therefore, there is a critical gap in the field of musculoskeletal modeling and simulation between solving for muscle forces with an explicit, but uncertain, objective function (or subset of them) and solving for the broad range of possible muscle forces that include muscle force combinations that are not feasible.

Bayesian inference methods are well suited for problems where we want to constrain the set of plausible solutions based on prior evidence and knowledge of the musculoskeletal system. This evidence could include information about physiology, measurement errors, and model-based uncertainties. Bayesian methods can use prior probability distributions based on previous knowledge of the system to help constrain the search space to within a set of reasonable bounds, and thereby estimate a plausible set of muscle forces. Markov Chain Monte Carlo (MCMC) algorithms sample from Bayesian models by searching through a multi-dimensional parameter space according to rules that compare proposed moves. The moves are evaluated based on a weighting function derived from a combination of a prior probability distribution and a likelihood function (36,37). Then, as the MCMC algorithm is iterated, the set of visited locations is used as a sample from the Bayesian posterior distribution of the unknown parameters (37,38). The output of an MCMC analysis is a sample from a posterior probability distribution, which numerically represents a set of equally plausible parameter vectors that could produce a result that is similar to the observed data.

Our aim is to evaluate the feasibility of using Bayesian inference methods to quantify the plausible range of muscle forces for human motion. For this initial feasibility assessment, we developed a prior based on a commonly used objective function (integrated cubed excitation), with uncertainty around a most likely value of that function, while keeping other musculoskeletal model parameters constant throughout the study (e.g., peak muscle forces, tendon slack lengths). Our prior was based on physiological hypotheses that muscle forces are distributed amongst agonist muscles and that co-activation of agonist and antagonist muscles is typically low for healthy human motions (22,39). We used an MCMC sampling algorithm in MATLAB and simulated an elbow flexion-extension task (reference motion) using OpenSim to explore the plausible excitations that could give rise to the reference joint trajectory. We then compared the excitations from MCMC to the known original simulation that generated the reference motion. Our aim for this paper is to present a workflow for performing MCMC analysis to sample plausible muscle forces for a measured motion with a musculoskeletal model. Ultimately, we hope that this workflow will allow movement scientists to appropriately characterize uncertainty in musculoskeletal simulations.

## Methods

### Musculoskeletal Model

We used an MCMC algorithm to identify a range of plausible muscle excitations that could produce an observed elbow flexion-extension motion in a simple musculoskeletal model. The musculoskeletal model was modified from a preexisting model (arm26; adapted from: (21)) available on the OpenSim 3.3 software package (Figure 1A; (9)). We further modified the arm26 model so that it had only a single mechanical degree-of-freedom (DOF), representing flexion and extension of the elbow joint. The elbow was actuated by six muscle-tendon units (Millard2012EquilibriumMuscle; (40)): three elbow flexors (brachialis, biceps brachii long head, biceps brachii short head) and three elbow extensors (triceps brachii lateral, triceps brachii medius, triceps brachii longus). We also modified the muscle-tendon properties so that the series elastic element for all six muscles were treated as rigid for computational speed.

**Figure 1.**
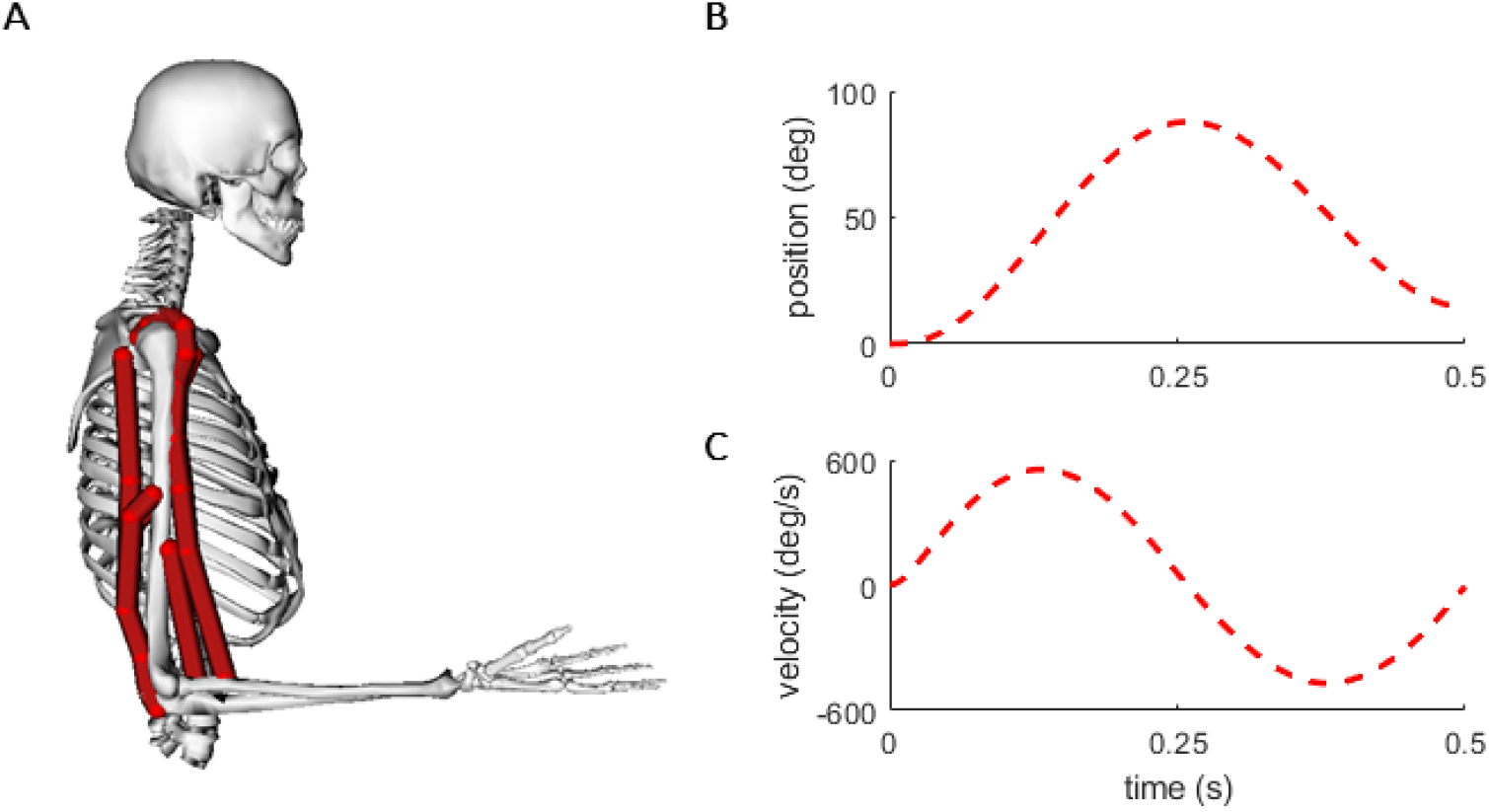
Elbow Musculoskeletal Model and Reference Data: A) OpenSim elbow model with the elbow posed at 90 degrees (1.57 rad; mid-point position). Red lines represent the paths for each of the six muscles in the model. The reference position (B) and velocity (C) trajectories used as input into the MCMC log likelihood function.

To create a unique “ground truth” reference motion to use for the MCMC analysis, we performed the following steps. First, we generated a smooth elbow flexion-extension trajectory that began at full elbow extension (0°), flexed to 90°, then extended back to full extension in a time period of 0.5 seconds. Next, we used this motion to generate a set of muscle excitations with the computed muscle control tool built into OpenSim (13). To account for the likelihood that human movements are not generated from perfect effort-optimal solutions as is assumed in computed muscle control, we then added noise to each of the muscle excitation trajectories (SD = 0.02 units of excitation), such that the excitations produced a small level of co-activation. We used this new set of muscle excitations in a forward simulation to produce a new elbow angle trajectory, which began with full elbow extension, flexed to about 90 degrees, then extended back to about 20 degrees (Figure 1B-C). This unique, non-effort-optimal trajectory was used as the reference motion for the MCMC analysis. The forces from this reference trajectory were used for comparison with the MCMC analysis described below.

### Markov-Chain Monte Carlo Analysis

We used an MCMC algorithm based on a Delayed-Rejection, Adaptive Metropolis (DRAM) formulation (36,41) to sample from the posterior distribution of parameters defining muscle excitations (Figure 2). To generate time-varying muscle excitation signals, we used a set of ten compact radial basis functions (CRBFs) that were summed over the time of the simulation (42). Each CRBF had a fixed center (c) and width (w) which were determined heuristically for this problem, while the amplitudes (A) of each CRBF were the parameters in the MCMC algorithm. For each muscle, the ten CRBFs were then summed:

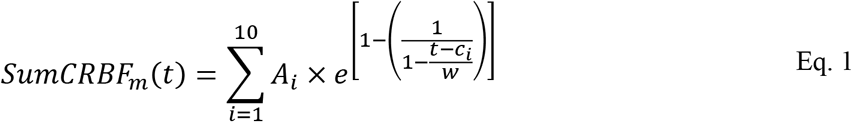

**Figure 2.**
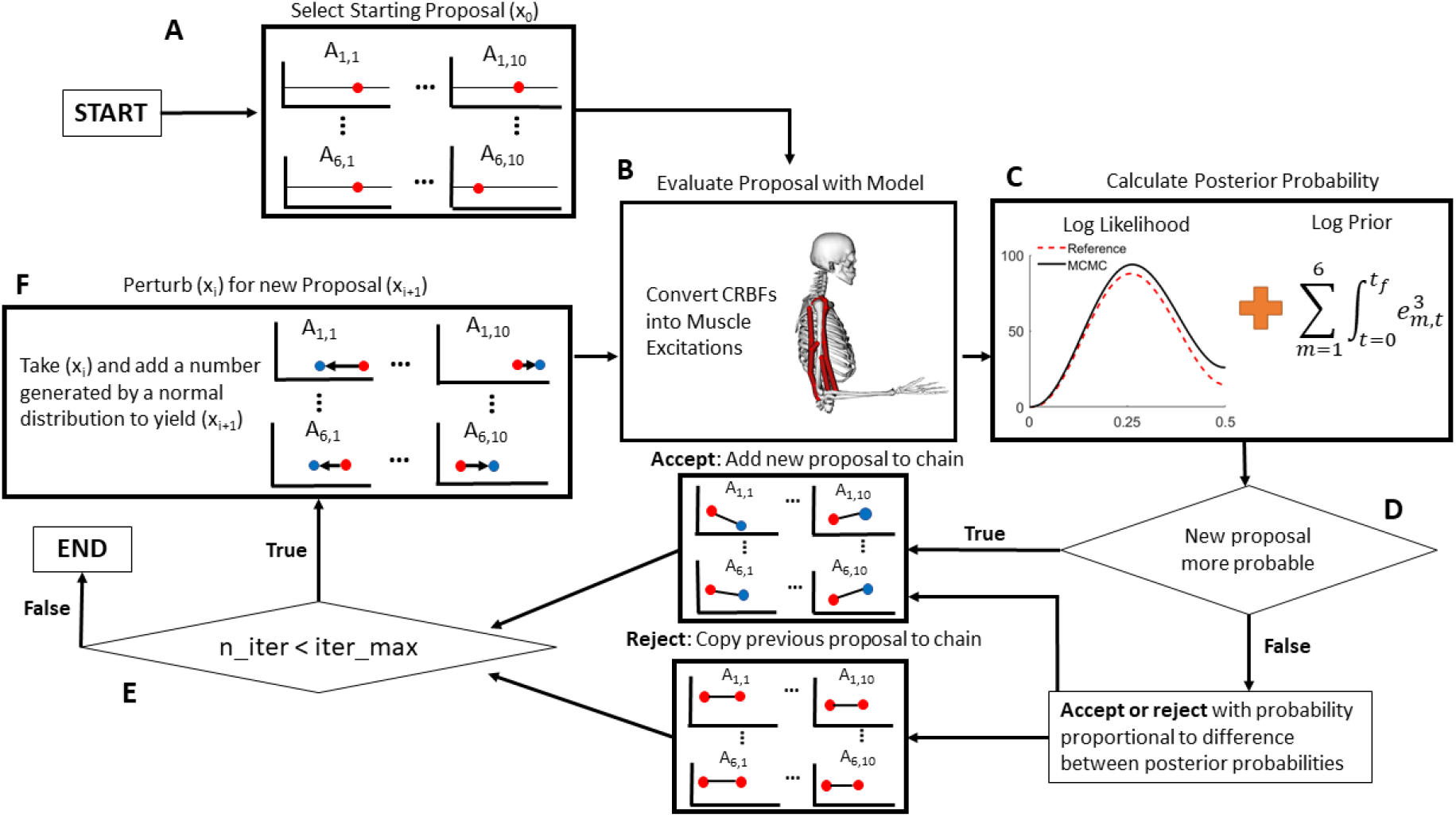
Flow Chart for MCMC and Elbow Flexion System: (A) The starting proposal for each parameter is drawn from a uniform distribution between [−5,0]. There are 60 parameters total representing amplitudes of the compact radial basis functions (CRBFs), 10 parameters for every muscle, where A_1,1_ is the amplitude of the first node of the first muscle, and A_6,10_ is the amplitude of the tenth node of the sixth muscle. (B) The proposal is converted from the set of CRBFs into a muscle excitations (Eqs. 1–2), which are given to OpenSim to generate a reference motion. (C) The posterior log-probability is calculated from the log likelihood (sum of square errors to the reference motion) and the log prior (the sum of muscle excitations (*e*) cubed). (D) The current proposal is accepted or rejected based on the change in posterior log probability from the original proposal to the new proposal (initial proposal is always accepted). (E) If the current iteration (n_iter) is equal to the pre-defined maximum iterations (iter_max), the MCMC exits, otherwise it generates a new proposal in (F) by perturbing the current proposal by a value drawn from a normal distribution. Further details on the algorithm and acceptance criteria are given in (36,41).

*A_i_* is the amplitude of the *i*th CRBF, and *t* is the time vector from 0 to final time (*t_f_* = 0.5 s). The centers (*c_i_*) were equally spaced between t = −0.05 and t=0.55 (intervals of 0.0667 seconds) and the widths (*w*) were equal to 0.0887 s, or 1.33 times the distance between consecutive centers. The *SumCRBF*s were then transformed such that they ranged from 0 and 1 using an inverse logit-transform (Figure 3). The inverse logit function was computed in a numerically stable way with Eq. 2, which ensures the exponential function is only applied to negative arguments. Each *A_i_* was further constrained to be between −30 and 30 (representing a uniform prior on individual coefficients), since values outside of these bounds, once converted with the logit transform, are approximately equal to 0 (for values less than −30) or 1 (for values greater than 30), therefore searching outside of these bounds becomes redundant.

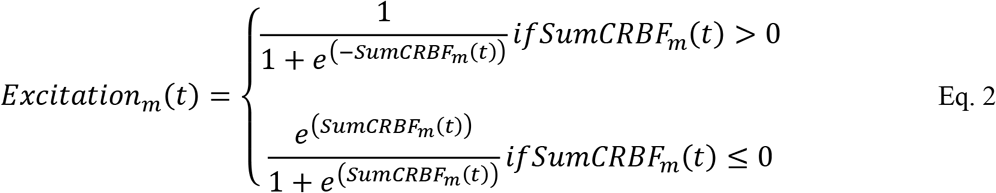

**Figure 3.**
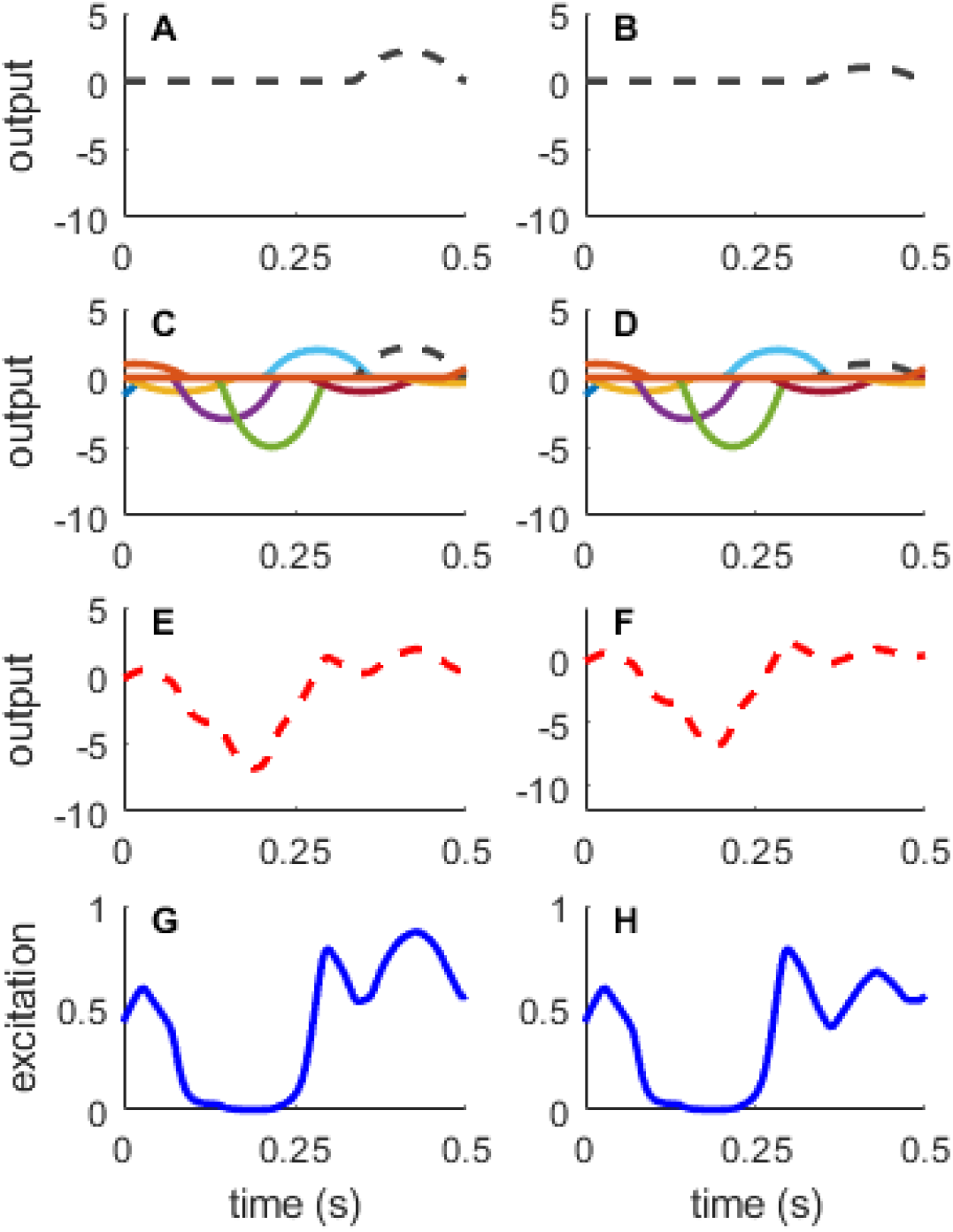
Generating muscle excitations from compact radial basis functions (CRBFs): The excitation signal for each muscle was determined by the amplitudes of ten CRBFs, each with a pre-defined center and width. (A) One CRBF shown individually. (C) All ten CRBFs with preset centers and widths, with varied amplitudes. (E) The sum across all ten CRBFs for a given muscle. (G) The summed output is then converted via an inverse-logit transform (Eq. 2) so that the excitation is constrained to be between 0 and 1. These excitations are then sent to the OpenSim model to perform the forward integration. The right column (B, D, F, and H) shows the effect of changing the amplitude of a single CRBF (black dashed line). Changing the amplitude of the CRBF affects the muscle excitation signal within the width of the CRBF, but the area outside of the CRBF remains constant.

The process for a standard Metropolis-Hastings MCMC algorithm begins by setting an initial, user-defined proposal for the amplitudes of the CRBFs, then simulates the model and compares the resulting output with the true data set. The parameters of the CRBFs define the set of muscle excitation trajectories that were used to perform a forward dynamics integration in OpenSim. The simulation used inputs from the muscle excitation trajectories and the initial states of the model (initial elbow angle (0°), angular velocity (0°/s), and initial muscle activations (all muscles = 0.05)), and calculated the motion of the model through a forward integration of the set of muscle excitations to generate the kinematics for the current iteration of the MCMC algorithm. The forward dynamics simulations account for the muscle excitation-activation and force-length-velocity relationships of the contractile elements. For each proposal, the simulated kinematics of the current proposal were compared to the reference motion by computing the sum of squares error between the reference and position and velocity trajectories of the proposal to generate a log likelihood function (Eq. 3).

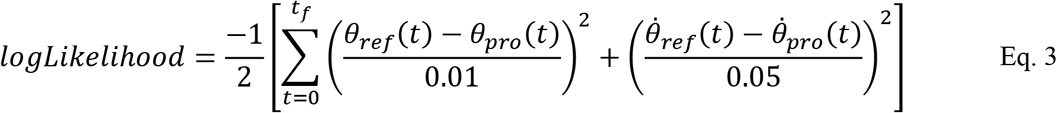

*θ*_*ref*_(*t*) is the reference angular trajectory, *θ*_*pro*_(*t*) is the position trajectory of the current proposal, 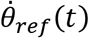 is the reference angular velocity trajectory, and 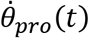 is the velocity trajectory of the current proposal. The denominators in each term were chosen to give each term a relatively equal contribution to the log likelihood function on the basis that a-priori all errors are of approximately equal contribution. In addition to the log likelihood function, the posterior probability also included a log prior function to represent the physiological hypothesis that the central nervous system distributes muscle forces across agonist muscles in a way that reduces muscle fatigue (22,39). The log prior function was designed such that the sum of the integrated muscle excitations cubed in the MCMC were similar to that of the sum of the integrated muscle excitations cubed from the reference data (Eq. 4).

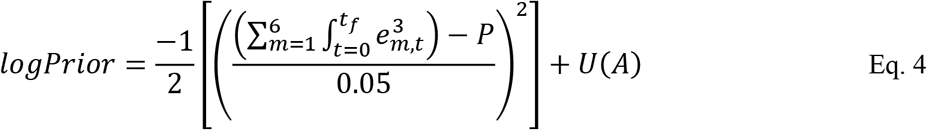

Here, *m* is an index corresponding to each of the 6 muscles, *t* is time and *e_m,t_* is the muscle excitation of the *m^th^* muscle and time *t*. *P* was equal to 0.07, which was the sum of integrated muscle excitations from the reference trajectory, and was used as a target center for the sum of the integrated muscle excitations cubed. The denominator for the log prior was selected to be arbitrarily wide, such that the initial search space was not overly constrained by the prior function definition. We note that with real experimental data, the *P* value used in Eq. 4 would not be directly measured, however users may compute a *P* value by running a static optimization (or related algorithm) with their experimental data, or a hierarchical prior over P can also be employed to express uncertainty in this value. Based on our constraints of the amplitudes of the CRBFs, the U function is 0 on the allowed range [−30,30] for each A value, and negative infinity outside that range.

After the user-defined initial proposal is simulated, a new proposal is generated by adding random noise to the previous proposal and then evaluating the new proposal with the model. New proposals are always accepted if the log posterior probability density (Eq. 5) increases compared to the previous proposal.

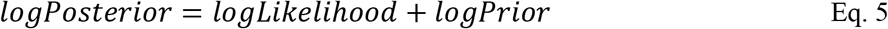

If the new proposal has decreased log probability density compared to the previous position, it is accepted or rejected with a probability based on the difference between the log probability density values of the two proposals. If the new proposal is accepted, it becomes the next sample in the chain. If the current proposal was rejected, we utilized a Delayed-Rejection algorithm to resolve rejected proposals. After a rejection of a proposal, instead of immediately retaining the previous proposal and advancing to the next iteration, a second-stage proposal is evaluated (36). The second-stage proposal is an intermediate point between the previous iteration in the chain and the rejected proposal (see (36) for mathematical details). If the second-stage proposal was also rejected, we copy the proposal from the previous iteration to the current iteration. Once this process is completed, a new proposal is generated and evaluated with the same methodology. This process continues iteratively until a predetermined stopping point is reached, usually based on the total number of iterations (Figure 2). Lastly, as the MCMC algorithm iterates, we used an Adaptive Metropolis algorithm to tune the proposal distribution using the history of the chain with the goal of obtaining reasonable results without excessive iterations (41). The Delayed-Rejection and Adaptive Metropolis algorithms work together to improve the overall efficiency of the Metropolis-Hastings algorithm without requiring the calculation of derivatives with respect to parameter values as needed in some other algorithms based on Hamiltonian Monte Carlo, for example.

Five parallel MCMC algorithms were run simultaneously (in parallel) starting from different initial proposals. The initial proposals for each of the sixty parameters, for each of the five parallel runs, were drawn at random from a uniform distribution between [−5,0]. The algorithm was set to terminate at 450,000 iterations, with the first 90,000 iterations discarded as burn-in (first 20%) before the results were evaluated.

### Performance Analysis

We evaluated the performance of the MCMC algorithm based on 1) the total time for the algorithm to run, 2) whether the algorithm converged, 3) the root mean squared error (RMSE) between the MCMC positions and velocities compared with the positions and velocities of the reference trajectory, and 4) whether the resulting muscle forces from the MCMC were comparable to the forces from the reference motion. The five parallel MCMC simulations were evaluated for convergence by calculating and plotting the rank-transformed values across all chains (43). When converged, these rank plots should show a uniform distribution between 0 and 1 in all chains. This indicates that each MCMC simulation is exploring the same space as the whole ensemble, as a subjective means of assessing convergence. If the rank plot is relatively flat across the range, then we declare that the MCMC algorithm has converged. In addition, we also evaluated the convergence diagnostics for the potential scale reduction statistic 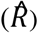 and effective sample size for each of the parameters (44,45). 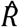 gives an indication of how well the parallel chains have mixed and reached equilibrium, with values less than 1.10 commonly accepted as converged. The effective sample size gives an estimate of the number of independent samples in the chain, since the MCMC chains will typically have some level of autocorrelation within a chain. Typically, an effective sample size of about 30 samples will allow for a reliable calculation of the means and standard deviations of the results according to the central limit theorem.

We also computed correlation coefficients between pairs of parameters to understand how parameters correlated across time (correlating amplitudes at each time node) or across muscles (correlating amplitudes for each muscle). We expected the correlation analysis to demonstrate the relationships inherent in the muscle redundancy problem. Therefore, we predicted that the forces generated by agonist muscles should have strong negative correlations with each other at a given time point (i.e., as one muscle increases force, an agonist muscle would decrease force to meet the torque requirements) while being positively correlated with forces in antagonist muscles (i.e., as one muscle increases force, an antagonist muscle would increase force to result in co-activations). Whereas we predicted the correlations within a muscle between adjacent time points (e.g., nodes 1 and 2) to have moderately strong negative correlations, since these parameters determine the excitation amplitude for overlapping regions of time (see Figure 3C and 3D), while correlations between distant time points (e.g., nodes 1 and 10) to have weaker correlations as these points represent vastly different aspects of the movement.

We compared the forces from the MCMC results to the force trajectories of each muscle from the reference motion. The forces from the reference motion were computed directly with OpenSim based on the muscle excitations used to drive the motion of the model to produce the reference position and velocity trajectories. For this analysis, we took 25 random draws from each of the five parallel chains, after the burn-in phase, and calculated the means and standard deviations of the position and velocity, and the muscle forces of the random draws, as well as the RMSE between the kinematics of the random draws and the reference motion. The MCMC algorithm was performed on an eight-core desktop computer with a 2.1 GHz Xeon-Silver 4110 processor and 32 GB RAM.

## Results

It took 71 hours for the MCMC algorithm to perform 450,000 iterations across the five parallel chains. The sampled trajectories tightly tracked the position (Average RMSE = 0.008° across the 125 random draws; Figure 4A) and velocity data (Average RMSE = 0.18°/s; Figure 4B). The sum of the integrated muscle excitations cubed in the posterior were approximately normally distributed as 0.15 ± 0.06 units of integrated muscle excitation cubed (Figure 4C). When evaluating the range of plausible muscle forces for each of the six muscles in the model, the three elbow flexors produced force at the beginning of the motion (Figure 4G-I) to accelerate the elbow into flexion. Then, the three elbow extensors decelerated the elbow during the mid-point of the motion (Figure 4D-F). Lastly, the elbow flexors muscles again acted at the end of the motion to move the elbow to its final configuration.

**Figure 4.**
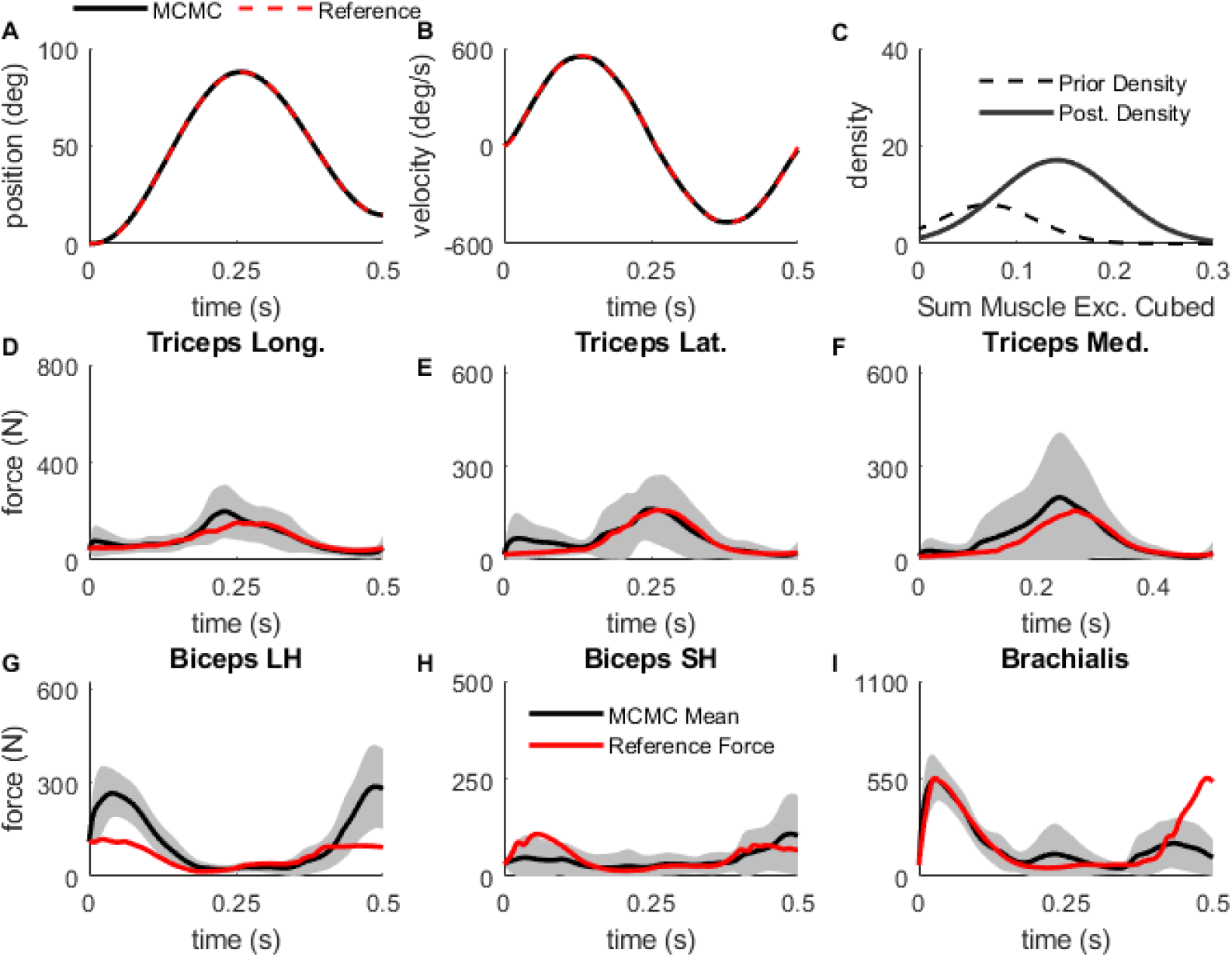
Random Draw Analysis: The position (A) and velocity (B) trajectories matched closely with the reference (red). (C) The prior and posterior (post.) density on sum of muscle excitations cubed. The posterior is shifted to center around slightly greater muscle excitations than the prior (0.09 ± 0.02). The mean and standard deviation of muscle force trajectories for triceps long head (D), triceps lateralis (E), triceps medialis (F), biceps long head (G), biceps short head (H), and brachialis (I) compared with the forces from the reference trajectory (red).

When evaluating the sampled force results against the forces that produced the reference motion, the average elbow extensor forces from MCMC matched extremely close with the reference. Overall, the average elbow flexor forces from MCMC matched well with the reference, although there were a few deviations. At the beginning of the motion, the biceps long head produced more force on average than the reference for that muscle, but the biceps short head produced less force than the reference. Then at the end of the motion, the brachialis produced less force than the reference, but the biceps long head produced more force than its reference. Overall, there was little evidence of any strong co-activation between agonist and antagonist muscles. The distribution of forces between agonist muscles was the primary difference between different MCMC draws, as is evident by the wider standard deviation range in muscle forces during the time that those muscles were active.

Overall, our rank plot analysis showed that the five chains were not fully mixed after the iterations ended. In some cases, the chains seemingly became stuck in local regions, for example chain 2 of node 3 in the triceps lateralis deviates from the other four chains (Figure 5A). The rank plots for each of the parameters did not have a flat distribution, indicating the parallel chains were not searching the same solution spaces (triceps lateralis; Figure 5C and 5D). Across all sixty parameters, the 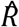 statistic ranged between 1.02 to 5.42, with only six of sixty parameters below the acceptable threshold of 1.10. The effective sample size was also low for the number of iterations performed, ranging between 12 and 50 samples (triceps lateralis; Table 1).

**Figure 5.**
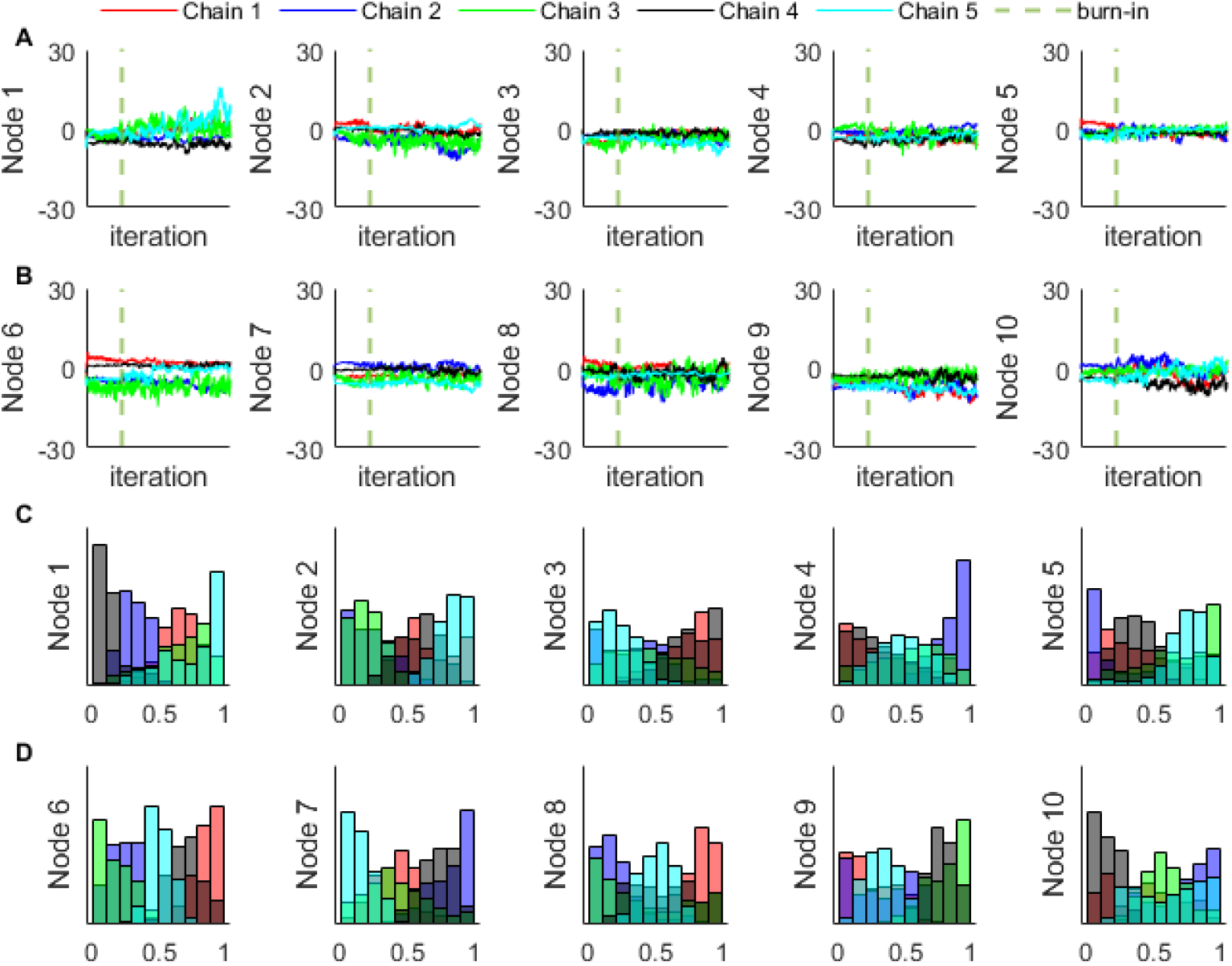
Chain and Rank Plots for the Triceps Lateralis: Chains (rows A and B) and rank plots (rows C and D) for each of the ten nodes for a representative muscle. Overall, the rank plots show that the four chains did not fully mix, although nodes 2, 3, and 9 have better mixing than nodes 1, 4, and 10.

**Table 1:**
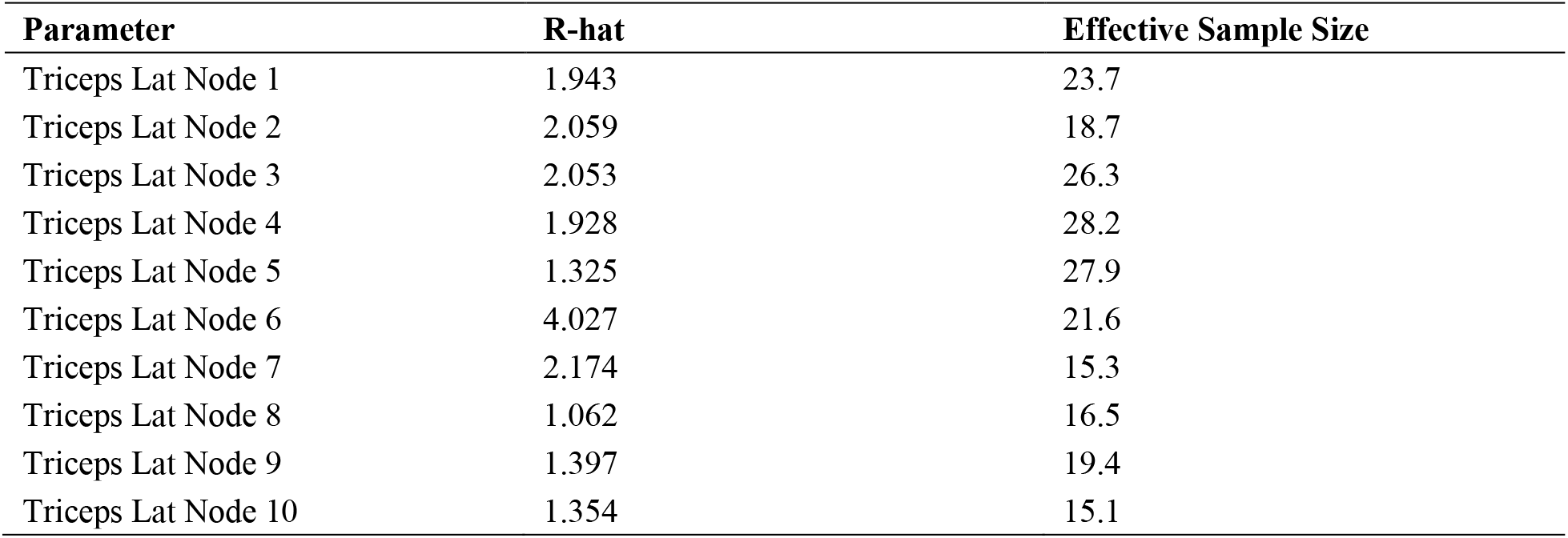
Convergence diagnostics for a representative muscle (triceps lateralis): the potential scale reduction statistic 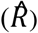 where values < 1.10 are considered to indicate that the chains are in equilibrium, and the effective sample size indicating the number of independent samples in the chain.

There were significant correlations between pairs of coefficients (Figure 6: between muscles for the first time node; all correlations statistically significant at p<0.001). For example, the negative correlation between the amplitude of the triceps long head and triceps medialis (r = −0.69) show that when the excitation is greater for one muscle, it is low for the other muscle. Other correlations reveal potential co-activation strategies explored by the algorithm, such as the positive correlation between brachialis and triceps medialis (r = 0.76). This correlation shows that when the excitation level is greater for the triceps medialis during initial elbow flexion, the brachialis excitation must be greater as well to match the kinematics of the movement at the time. The negative correlations for agonist muscles and positive correlations for antagonist muscles seen here fit our predictions, however there were a few cases where agonists muscles had negative correlations or antagonists had positive correlations (e.g., brachialis and triceps long head, r = −0.79). Additionally, there were also significant correlations between nodes of each muscle (Figure 7; all correlations statistically significant at p<0.001). Generally, correlations between adjacent time point nodes (the plots along the diagonal of Figure 7) had negative correlations, as we expected. However, there were other non-intuitive results from the correlation analysis, as some of the distant time nodes had stronger correlations that we predicted (e.g., node 2 and 10 of triceps long head; r = 0.60).

**Figure 6.**
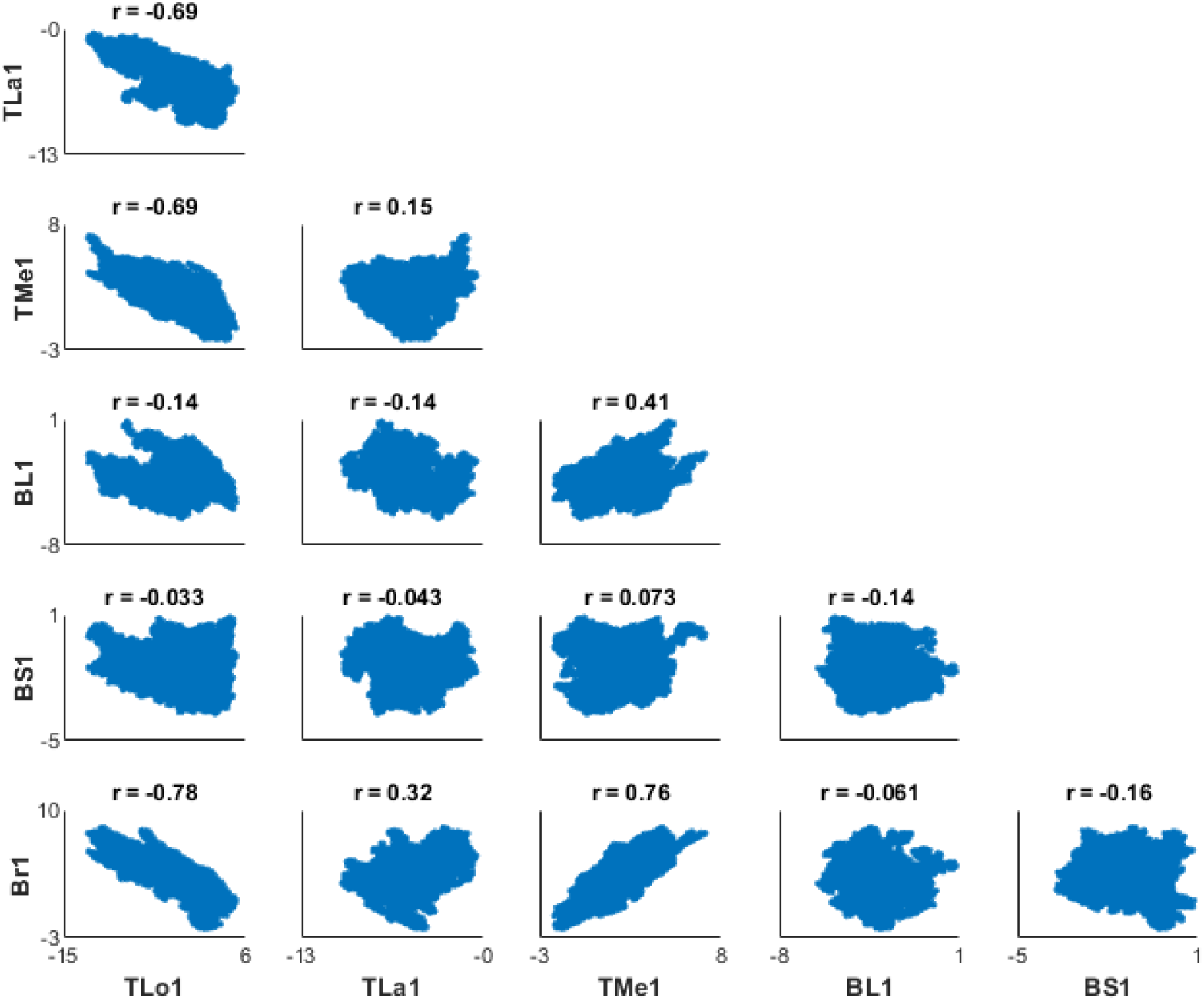
Node 1 Correlations: Correlation plot between the six muscles in the model for the parameters representing the first time point, or node: triceps long head (TLo1), triceps lateralis (TLa), triceps medialis (TMe1), biceps long head (BL1), biceps short head (BS1), and brachialis (Br1). All correlations are statistically significant at p < 0.001.

**Figure 7.**
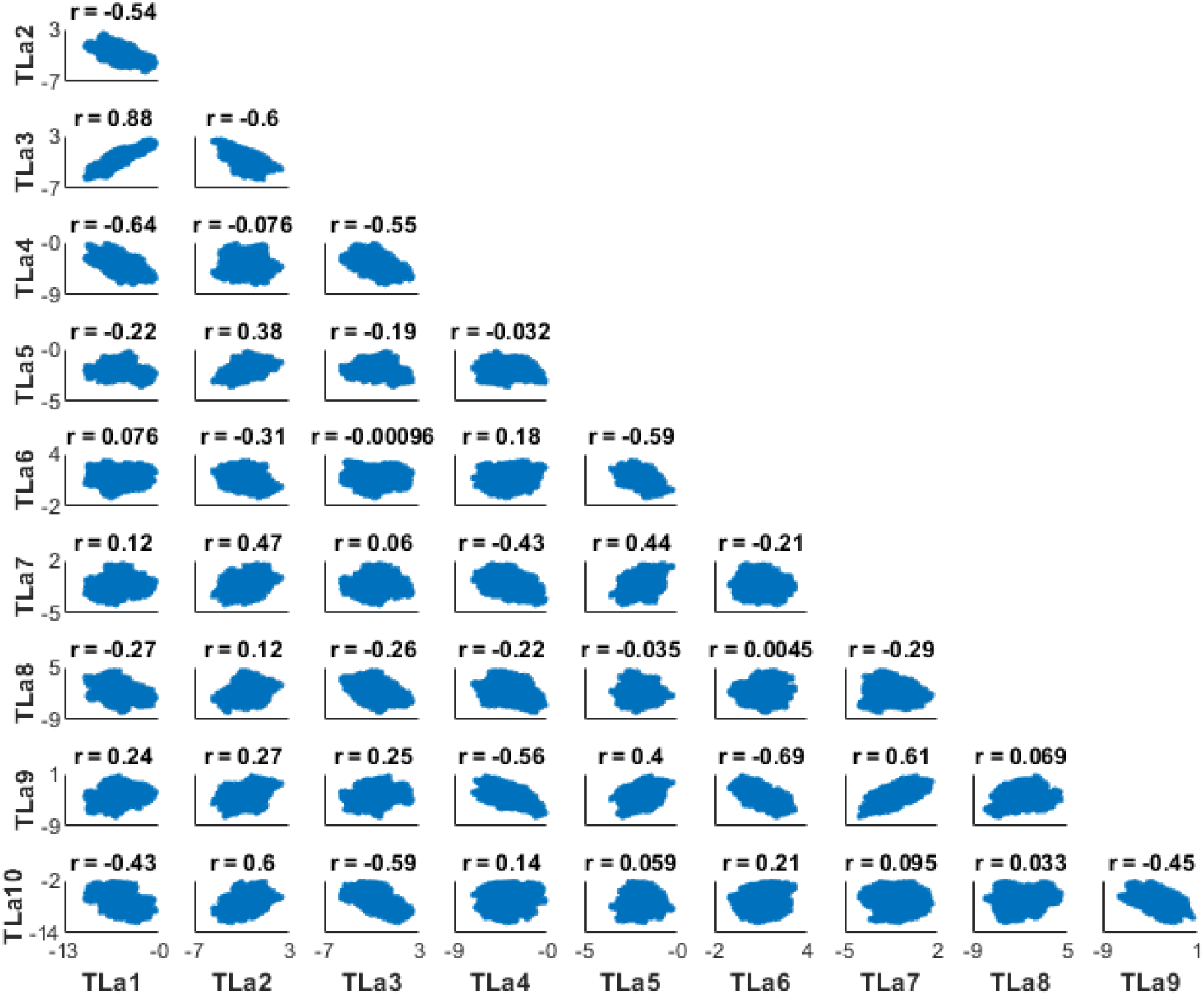
Triceps Lateralis Correlations: Correlation plot between the ten nodes of the triceps lateralis muscle. All correlations are statistically significant at p < 0.001.

## Discussion

A key question in motor control and biomechanics is how our nervous system controls our muscles during movements, and therefore how muscle forces are distributed across muscles (8,16,22). Musculoskeletal modeling and simulation present a way to estimate the muscle forces during a given motion, but these methods rely on several modeling assumptions. Standard methods for estimating muscle forces result in estimates of a single time-series of muscle forces for a given movement, which disguises the real uncertainty that accompanies the unknown model parameters and other related assumptions used for musculoskeletal modeling studies. In this paper, we presented a workflow and evaluated the feasibility of using Bayesian estimation and MCMC to find a plausible set of muscle forces that could result in an observed trajectory in a simulated elbow flexion-extension task, while potentially allowing for any mathematical hypothesis about the motor control problem. In order to define the potential solution space, we used a prior probability distribution intended to represent physiological evidence that muscle forces are distributed across agonist muscles in a way the reduces muscular effort (22). This approach resulted in a range of muscle forces that reproduced reference kinematics and generally matched with muscle forces from the reference trajectory (i.e., the ground truth), while illustrating the array of plausible combinations of muscle forces that could give rise to the reference elbow motion.

Our Bayesian approach allowed for uncertainty in the kinematic data through the use of a likelihood function that represented error in the positions and velocities from the reference trajectory. Our results had little variability in the kinematics relative to the reference trajectories (Figure 4A and 4B), but future implementations could allow for greater uncertainty in these trajectories by increasing the denominator in each of the terms of the likelihood function (Eq. 3), especially if there was increased uncertainty in the collected data. For example, if kinematic data were collected via retroreflective markers with greater likelihood for measurement error, due to a greater probability of skin or clothing movement relative to bony landmarks, the MCMC implementation could set wider distributions for those variables to capture the uncertainty in the data.

Our approach builds upon previous work by utilizing a prior probability function, which is based on the physiological hypothesis of minimization of muscle effort, to constrain the solution space and provide more plausible bounds on minimum and maximum forces during the movement. Representing the hypothesis of minimization of muscle effort with a prior probability function allows for a balance between the singular solution from standard optimization routines and the relatively unbounded solutions from previous work (29–31). Computing the sum of the cubed muscle excitations integrated over time is one hypothesis for how the nervous system deals with the muscle redundancy problem during motor control, and it is a common objective function used in many musculoskeletal modeling optimizations (8). Our results illustrate that there are many different combinations of muscle force trajectories, with comparable total muscle effort, that can give rise to very similar kinematics. One potential advantage of using this prior probability function is that it can allow the user to change the parameters of the function to specify a broad set of plausible motor control hypotheses, thereby shaping the range of the plausible muscle forces. Other mathematical forms are also easily possible. For example, creating a narrower prior probability density, which would indicate a high degree of confidence in the hypothesis of minimizing muscle effort, would shrink the width of the shaded standard deviations in our results (Figure 4). While our study focuses on the uncertainty in kinematic data and model of motor control (i.e., the objective function), a potential advantage of this Bayesian approach is the ability to include many different sources of data or other hypotheses within the prior density function (46,47), or include other types of uncertainty such as model parameter estimation or different types of measurement error (17–20,26–28). For example, future implementations of this paradigm could utilize EMG data for some muscles, while still constraining the solution space for muscles without EMG data (expanding on (29)).

We expected our correlation analysis to reveal clear demonstrations about how the nervous system can solve the muscle redundancy problem, with negative correlations between agonist muscles and positive correlations between antagonist muscles at a given time point. We further expected that correlations within muscles across time points would reveal negative correlations between adjacent nodes and flatter correlations between distant nodes. While many of the correlations followed our predictions, several correlations revealed unexpected relationships between parameters, suggesting that there may be non-intuitive relationships in how the muscle redundancy problem gets solved for a given motion. This is yet another advantage of the Bayesian approach, revealing potentially non-intuitive correlations between parameters in a system.

Other previous work has used sensitivity analyses for musculoskeletal modeling studies by systematically varying the objective function and re-solving the optimization problem for each function (16,23–25) or searching through a range of potential unknown modeling parameters (17,26–28). While these approaches can allow for an exploration of the sensitivity of modeling assumptions to the overall results, there can be important limitations to traditional sensitivity analyses. Typical sensitivity analyses vary model parameters at random based on known or predicted measurement error for a set of parameters of interest (e.g., muscle insertion coordinates, or tendon slack lengths). The limitation with this approach is that it could end up evaluating sets of parameters that are highly unlikely to produce the measured data. A Bayesian inference approach differs from these traditional sensitivity analyses as the MCMC algorithm will only search through spaces where parameter values give rise to reasonable predictions of the data. For example, the MCMC algorithm will reject proposals with extremely low probability and often accept proposals with high probability. Therefore, the computational time spent in regions of low probability is minimized, while the algorithm explores the regions of high probability. This can dramatically improve computational efficiency when exploring complex search spaces.

One potential obstacle in implementing MCMC with time-varying signals is that the problem has the potential to be extremely high-dimensional. For example, if each hundredth of a second was represented with an amplitude for the 0.5 second simulation, we would have to search 300 total parameters (50 nodes times six muscles). We were able to reduce the dimensionality of the problem by using CRBFs to compose the muscle excitation signals used in our forward simulations. CRBFs are useful for generating time-varying signals for MCMC problems as they allow complex time-series trajectories to be represented with a few parameters (42). CRBFs also have an advantage over other potential basis functions such as Chebyshev polynomials or Fourier series because they decouple the parameters from each other. For example, in a Chebyshev polynomial series, changing the value of just one coefficient will change the entire time-series trajectory, sometimes in dramatic ways (see Additional Files; Figure A1). Instead, the parameters (representing amplitudes) of the CRBFs change the shape of the time-series trajectory only within the preset width around the center of the individual function (Figure 4).

## Limitations

While this approach shows promise by producing muscle forces that compare favorably with the ground truth without being bound to singular solutions, there are still improvements that need to be made. Our rank plot analysis and convergence diagnostics 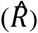 show that our MCMC chains were not converged, indicating that each of the parallel chains were not searching the same solution space throughout the MCMC. The lack of convergence is a limiting factor to our methodology now, and one that is bound by current computational speed and MCMC algorithm efficiency. To verify that the algorithm we chose would converge when used for simpler mechanical models, we used it to successfully recover model parameters of a mass-spring-damper system (see Additional Files). Improvements in algorithmic efficiency are critical to make this approach feasible for musculoskeletal modeling studies, especially studies that use more complex musculoskeletal models (e.g., models for gait). Future work in this area requires developing computational techniques able to search high dimensional spaces in an efficient manner. Once greater algorithmic efficiency is achieved, more complex musculoskeletal models could be included in this type of analysis (e.g., gait models) and more parameters could be included (e.g., muscle model parameters like peak isometric muscle force or tendon slack length). An alternative approach could include using differentiable musculoskeletal models or using algorithmic differentiation for OpenSim models (48), but each of these approaches also requires considerable computational set up time and expertise, which creates issues for broad applicability and reproducibility. Using a differentiable musculoskeletal model would allow for some different choices in MCMC sampling algorithms, such as algorithms that can utilize the slope of the solution space to better identify efficient ways to move with each iteration, and therefore converge in a faster amount of time.

## Conclusion

Altogether, this paper illustrates the type of analyses we could achieve with a Bayesian inference method applied to musculoskeletal modeling and simulation, allowing for the estimation of plausible muscle forces that could have generated an observed set of kinematics. The output from this analysis could then be applied to other outcome measures relevant to biomechanical modeling, such as providing plausible values for metabolic energy or joint contact forces in musculoskeletal models for a given motion. This approach can allow us to better quantify the uncertainty that is characteristic to musculoskeletal modeling and simulation studies. Improvements to the algorithmic efficiency will make this method more feasible to merge into a new standard of musculoskeletal modeling and simulation. Therefore, further developments for MCMC search algorithms which specifically are usable for common musculoskeletal simulation tools (or other problems without easy access to model derivatives) should be at the forefront of future research, since these kinds of mechanics problems are difficult to sample from. Doing so will allow researchers to better represent uncertainty in musculoskeletal modeling and simulation studies in the future.

## Declarations

### Ethics approval and consent to participate

Not applicable

### Consent for publication

Not applicable

### Availability of data and materials

Code is available at: https://github.com/russelljohnson95/MSM_Modeling_and_MCMC

### Authors’ contributions

Study concept and design (RJ, DL, JF), algorithm implementation and data analysis (RJ, DL), preparation of the manuscript (RJ), critical review and revision of the manuscript (RJ, DL, JF). All authors read and approved the final manuscript

## Acknowledgements

Not applicable

