## Additional Material for "Using Bayesian Inference to Estimate Plausible Muscle Forces in Musculoskeletal Models"

Additional Files

*MCMC Feasibility with Musculoskeletal Models using Chebyshev Polynomials*

One potential way to generate complex time series trajectories is with the use of orthogonal basis expansions such as Fourier series or Chebyshev polynomials. These can be thought of as a sequence of multiple individual basis functions such as sine or cosine functions that are summed together to generate a trajectory. Generating complex time series excitations, which are then sent to the OpenSim forward integration, was necessary for our MCMC approach with the musculoskeletal model. During our initial development of the MCMC paradigm with the musculoskeletal model, we used Chebyshev polynomials to generate our time series excitations for each muscle. To generate complex trajectories to control each muscle, we needed to include multiple (e.g., 10) coefficients for each Chebyshev polynomial function. Each Chebyshev polynomial is generated through the generalized form (with the Rodrigues formula; Eq. A1), multiplied by an amplitude coefficient (free parameter), and summed across the ten basis functions (Eq. A2).

$$T_n(x) = \frac{(-2)^n n!}{(2n)!}\sqrt{(1-x^2)}\,\frac{d^n}{dx^n}(1-x^2)^{n-\left(\frac{1}{2}\right)} \qquad \text{Eq. A1}$$

$$\mathrm{n} = [0{:}9],\ \mathrm{x} = [-1,1]$$

$$SumCheby_m(x) = \sum_{n=0}^{9} A_n \times T_n(x) \qquad \text{Eq. A2}$$

where $T_n(x)$ is a single Chebyshev basis function, $n$ ranges from 0 to 9 to give ten total basis functions for each muscle in our problem, $x$ is a continuous value that ranges from -1 to 1, and $A_n$ is the amplitude of the $n$th Chebyshev basis function. The $SumCheby_m(x)$ can then be converted via the inverse logit transform to constrain the muscle excitation to the range [0,1] (Eq. 2) and finally mapped from $x$ = [-1,1] to the time range of the simulation (e.g., $t$ = [0, 0.5]).

However, although orthogonal basis expansions are extremely efficient in deterministic function fitting, these types of functions can become an issue for Bayesian sampling problems, because changing one coefficient for a trajectory will change the output of the whole trajectory in time. Therefore, with many different parameters, each affecting the output along the whole time series, the parameters can become highly correlated which makes it difficult for the MCMC algorithm to move efficiently through the space. To illustrate this issue, here we present how a muscle excitation can drastically change across time, with the modification of one coefficient (Figure A1).

To address these issues, we then changed our process for generating muscle excitations to that of compact radial basis functions, where changes to a single coefficient only affect the trajectory within a certain compact region in time, effectively decorrelating widely separated coefficients.

*Mass-Spring-Damper Model*

**Methods:**

We evaluated the performance of the DRAM MCMC algorithm for estimating the parameters of a simple mass-spring-damper system for a simulated trajectory as a test case for inference on dynamical systems. We focused on attempting to recover the known set of parameters in a simple mechanical system as a precursor to using Bayesian methods on a more complicated mechanical system. The mass-spring-damper system consisted of a mass connected to the ground via a nonlinear spring with a variable stiffness in parallel with a damper (Figure A2). The variable spring stiffness had three parameters that defined its force-displacement relationship: two stiffness coefficients ($k_1$ and $k_2$) and the threshold ($T$) which defined when the spring stiffness transitioned between $k_1$ and $k_2$. The damper was defined with a single coefficient ($c$).

The mass-spring-damper system was used to generate a reference motion (Figure A2) of the mass based on specifically chosen values of the five parameters defining the system and the initial position ($x_0$) and initial velocity ($\dot{x}_0$) of the mass (Table A1). The motion of the block was simulated for 20 seconds. After the motion was simulated, we added noise to the trajectory from a Gaussian distribution with a standard deviation of 0.005 m which represented some realistic measurement error. We repeated the simulation to generate four total trials, with noise added for each trial. For the mass-spring-damper system, each new proposal during the MCMC iterations were evaluated based on the log posterior density of the resulting simulated trajectory. The log posterior density is calculated from the sum of squared errors calculated between the reference position trajectory for the four reference trials and the motion generated by the current proposal's parameters (the log-likelihood) plus the log of the prior probability density over the parameters.

We ran the same DRAM MCMC algorithm as for the musculoskeletal model (Figure A3), and we set the MCMC to run for 30,000 iterations, with the first 15,000 iterations considered "burn-in," which are discarded before the final analyses. The number of iterations for the MCMC algorithm was determined heuristically by evaluating the trade-off between convergence diagnostics and computational time. The burn-in is used primarily as a means of enabling convergence of the algorithm to the high-probability region before beginning analyses.

Five separate MCMC simulations were performed in parallel, each began from a different set of initial proposals. For each simulation, the prior probability density was created for each parameter by generating a wide gaussian distribution with the center defined by adding noise to the real parameter value. From there, the initial proposals were drawn from the prior probability density for the given MCMC simulation. We assessed convergence using a rank plot analysis to assess whether each chain was exploring similar solution spaces throughout the 15,000 iterations after the burn-in phase (1). We also calculated the potential scale reduction statistic ($\hat{R}$) and effective sample size for each of the seven parameters (2,3). After convergence was assessed, we then used a random draw analysis to evaluate how well the position trajectories from the MCMC analysis matched with the reference trajectories. To perform the random draw analysis, we selected 20 samples from the five chains (after burn-in) and calculated the position trajectory that would result from the parameters for each sample. This method enabled us to evaluate goodness of fit in a simple way without storing the entire trajectory for each of the 15,000 iterations.

The performance of the MCMC algorithm with the mass-spring-damper system was evaluated based on two main criteria: 1) The ability of the algorithm to find the parameters of the system that reproduced the reference motion and 2) the computational time necessary for MCMC convergence.

**Results:**

The MCMC algorithm performed 30,000 iterations across five parallel chains in 40 minutes. The results from the MCMC were centered around the true values of the parameters for each of the five runs. The five parallel MCMC simulations were considered to have converged since the rank plots for each parameter were relatively flat across the distribution (Figure A4D), each parameter had a $\hat{R}$ that was less than 1.02 (Table A1), and an effective sample size over 200 (Table A1). The random draw analysis showed that the positions from each of the drawn samples matched closely with the positions from the reference trials (Figure A4E).

Some of the parameters, such as the mass and damping coefficient, were determined to be correlated within the model (Figure A5). These correlations are not surprising in dynamical systems. Uncovering these correlations is an advantage of the MCMC approach as we discover the full complexity of the inference problem and can better assess the information we have about the parameter values. Overall, the MCMC algorithm performed well, recovered the correct parameter values, and provided a realistic assessment of the full remaining uncertainty in those values with a reasonable computational effort.

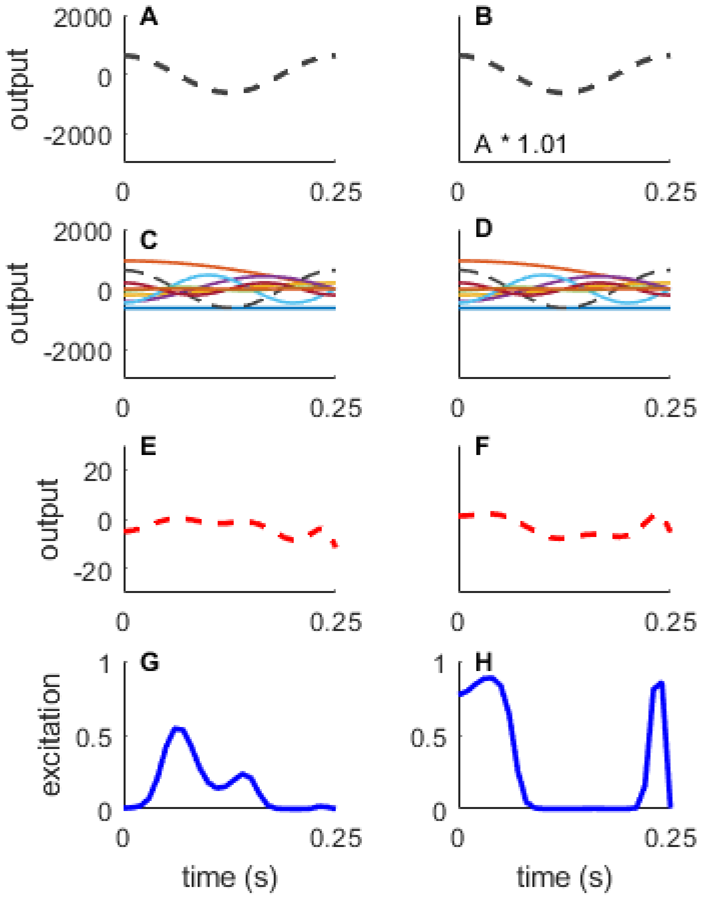


**Figure A1. Generating muscle excitations using Chebyshev polynomials:** A selected portion of the Chebyshev polynomials to illustrate the sensitivity of the time series muscle excitations to a single parameter (A) One individual polynomial (see Eq. A1) (C) All ten polynomials overlayed (E) The sum across all ten polynomials for a muscle. (G) The summed output is then converted via an inverse-logit transform (Eq. 2) so that the excitation is constrained to be between 0 and 1. The right column (B, D, F, and H) shows the effect of changing the amplitude of a single polynomial (black dashed line). The new polynomial was generated by multiplying the original signal of the $5^{th}$ polynomial by 1.01. In this case, a small change in the amplitude of a single polynomial can result in very different excitation signals across all time points (compare results of G and H). This sensitivity is why it is difficult for the MCMC algorithm to handle using Chebyshev polynomials to generate the muscle excitation signals.

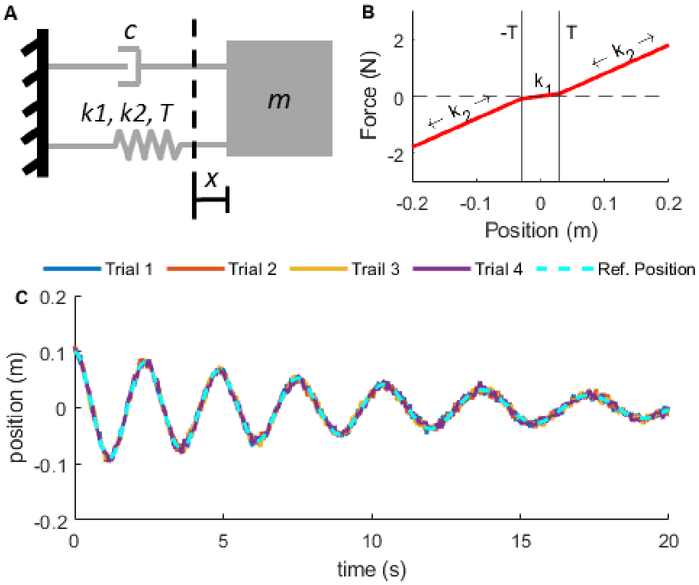


**Figure A2: Schematic of mass-spring-damper system** (A) The mass-spring-damper system consisted of a mass (*m*), a damper (*c*), and a nonlinear spring with stiffnesses of $k_1$ or $k_2$, with *R* being equal to the threshold to switch between $k_1$ and $k_2$ (*R* = 0.02 m), while *x* shows the displacement of the mass with respect to the slack length of the spring. (B) Force-displacement relationship for the variable stiffness spring, where the threshold (*R)* is indicated by the vertical solid lines. The spring stiffness $k_1$ occurs between -0.02 and 0.02 m, while $k_2$ occurs outside of those bounds. (C) Simulated trajectories of the model with added noise for the four trials (see Table 1 for model parameters).

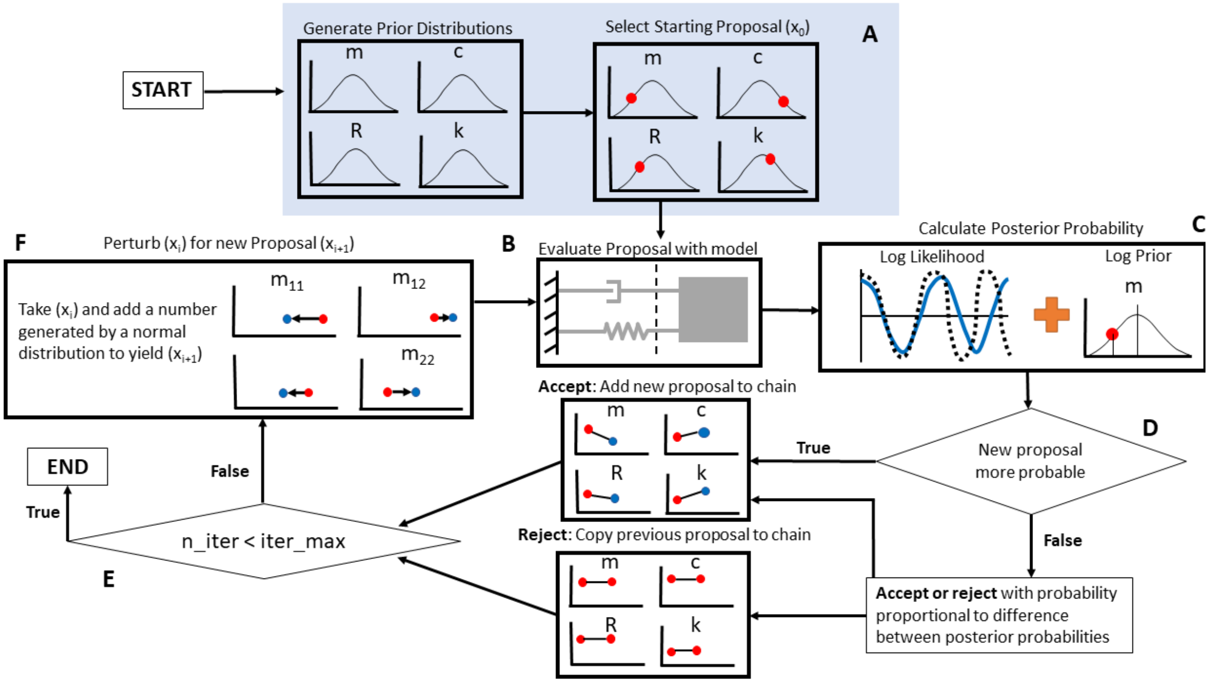


**Figure A3: MCMC flow chart with mass-spring-damper system** (A) Each simulation begins by sampling from the prior distribution to get the initial proposal (red dots in blue inset). (B) The initial proposal values are then used to simulate the dynamics of the system with those parameters. (C) Then, the log-posterior-probability is calculated by adding the log likelihood (sum of square error in position trajectory) plus the log prior log-density of the proposal (log(p(data | params)) + log(p(params)). (D) The current proposal is accepted or rejected based on the change in posterior log probability from the original proposal to the new proposal (initial proposal is always accepted) see text for details on how proposals are evaluated. (E) Check to see if the number of iterations (n_iter) is equal to the pre-determined iteration maximum (iter_max): if true, end the simulations. Otherwise, the sequence will continue with (F) Generate a new proposal as a perturbation from the current proposal.

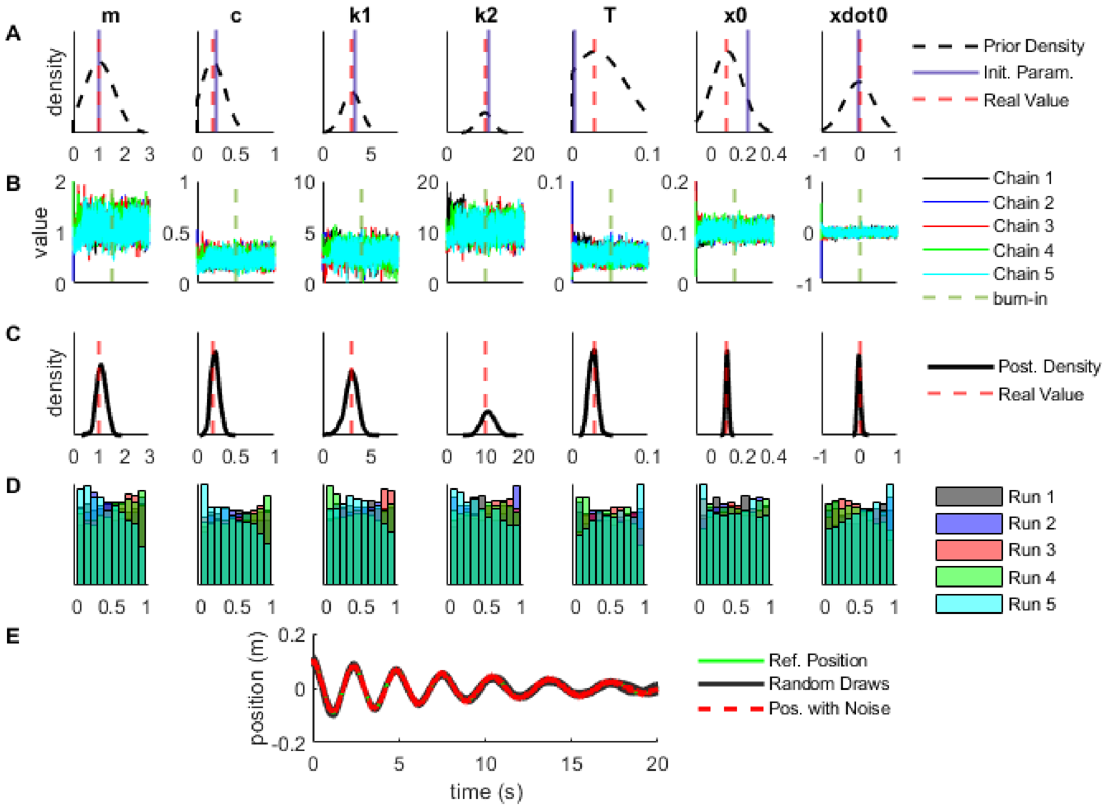


**Figure A4. Random Draw Analysis:** (A) The prior probability density, along with the real values and initial proposals (Table 1). (B) The random walk chain for each of the seven parameters from a representative run, where the vertical dashed line marks the end of the burn-in phase. (C) The posterior marginal densities along with the real value. (D) The rank plots generated for five separate MCMC simulations. This demonstrates convergence of the algorithm. (E) The results from 20 random draws from the chain, overlayed with the reference position data with added noise (from one representative trial) are shown.

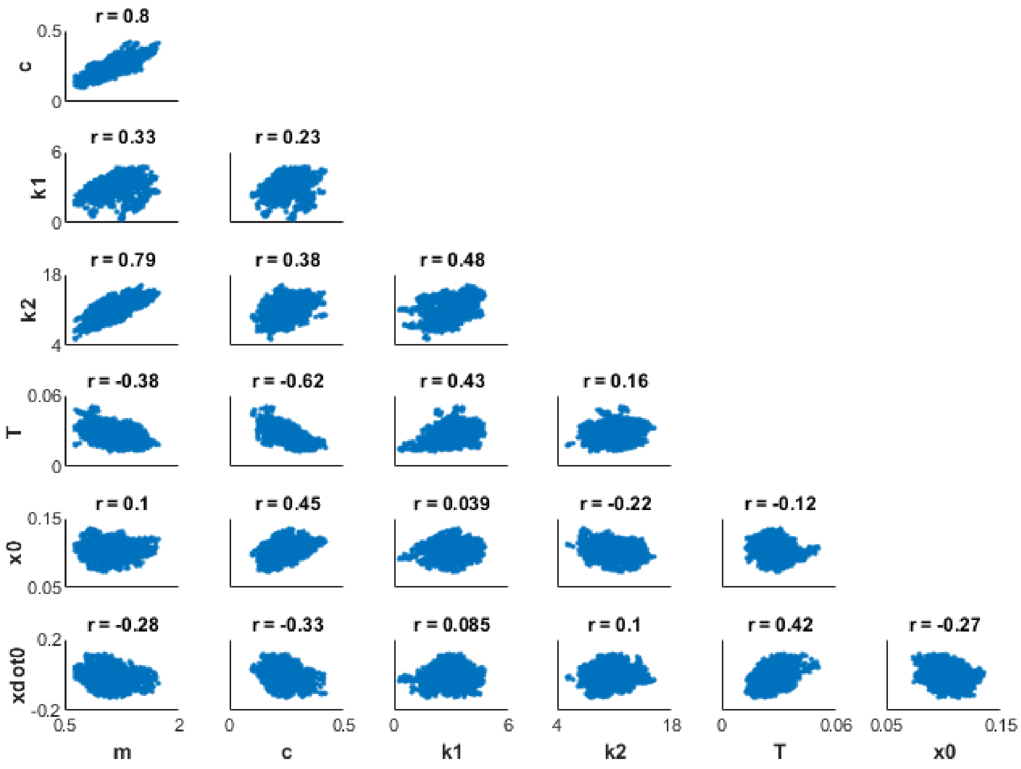


**Figure A5. Correlations between Parameters:** It is typical in mechanics problems to see correlations between the inferences for various variables in the system. Here, each subplot shows the correlation between two variables along with the computed correlation coefficient. For example, there is considerable correlation (R = 0.80) between the mass value (m) and the damper coefficient (c) such that the parameters remain plausible provided that mass and damper coefficient increase or decrease together. Whereas there is approximately no correlation between the initial position ($x_0$) and the stiffness 1 parameter (k1) (R = 0.04).

**Table A1:** The real values for each of the seven parameters of the mass-spring-damper system, the posterior mean and standard deviation (SD) from the MCMC result. Convergence diagnostics: the potential scale reduction statistic ($\hat{R}$ where values < 1.10 are considered to indicate that the chains are in equilibrium, and the effective sample size indicating the number of independent samples in the chain.

| **Parameter** | **Real Value** | **MCMC Posterior Mean ± SD** | **$\hat{R}$ statistic** | **Effective Sample Size** |
|---|---|---|---|---|
| Mass (kg) | 1.00 | 1.096 ± 0.193 | 1.009 | 249 |
| Damping (N s/m) | 0.20 | 0.231 ± 0.052 | 1.004 | 231 |
| Stiffness 1 (N/m) | 3.0 | 3.071 ± 0.577 | 1.009 | 259 |
| Stiffness 2 (N/m) | 10.0 | 10.63 ± 1.68 | 1.006 | 345 |
| Stiffness Threshold (m) | 0.03 | 0.029 ± 0.007 | 1.013 | 283 |
| Initial Position (m) | 0.10 | 0.101 ± 0.009 | 1.001 | 280 |
| Initial Velocity (m/s) | 0.00 | -0.008 ± 0.042 | 1.010 | 303 |

99
